# Rac1 inhibition slows down the segmentation clock and increases the somite size

**DOI:** 10.1101/2025.09.11.675517

**Authors:** Maria Pappa, Charisios D. Tsiairis

## Abstract

The presomitic mesoderm (PSM) is a transient embryonic tissue patterned along the anterior-posterior (A/P) axis of vertebrates that periodically generates somites at its anterior end. This process, termed somitogenesis, is driven by waves of gene expression that travel from the posterior to the anterior end of the PSM under the contol of the segmentation clock, a molecular oscillator. Using a primary culture system, we performed a small-scale chemical screen targeting cellular mechanical properties and we identified Rac1, a small GTPase, as a key regulator. Inhibition of Rac1 markedly altered the periodicity of gene expression waves and disrupted somite spatial organization. Further analyses showed that Rac1 inhibition modified gene expression patterns, notably affecting markers of the Wnt and Notch pathways which are central regulators of segmentation. These findings reveal a previously unrecognized role for Rac1-mediated signaling in coordinating the mechanical and molecular aspects of segmentation dynamics.

## Introduction

Somitogenesis is a fundamental morphogenetic event in early vertebrate embryogenesis that occurswithin the presomitic mesoderm (PSM), a transient mesenchymal tissue. This process ptoduces somites, which are epithelial structures that form periodically and later give rise to vertebrae, skeletal muscle, and dermis (Pourquié, 2001). Somite are generated sequentially, driven by waves of gene expression that propagate from the posterior to the anterior PSM. Each time a wave reaches the anterior boundary, a new pair of somites is formed. These waves are kinematic in nature, emerging from coordinated oscillatory gene expression at the single-cell level (Lauschke et al., 2013). The oscillations primarily involve and are regulated by the Notch, Wnt, and Fgf signaling pathways, organized in a molecular network known as the segmentation clock (Dequéant et al., 2006; Goldbeter and Pourquié, 2008). Beyond synchronized oscillations, spatial differences in oscillation frequency and phase across the PSM (Lauschke et al., 2013; Tsiairis and Aulehla, 2016) are essential for shaping wave patterns and establishing anterior– posterior (A/P) polarity, enabling cells to acquire distinct positional identities along the axis.

PSM cell identity and behavior are tightly regulated by graded signaling of Wnt, Fgf, and retinoic acid (RA) (Aulehla and Pourquié, 2010). High Fgf and Wnt activity in the posterior PSM maintains cells in an undifferentiated, mesenchymal state, supporting elevated motility and rapid oscillations (Aulehla and Herrmann, 2004; Dubrulle and Pourquié, 2004). Gradients of Fgf8 and Wnt3a, shaped by transcriptional regulation and mRNA decay, activate downstream targets such as Dusp4, Brachyury (T), Tbx6, and Msgn1 (Aulehla et al., 2003; Dequéant et al., 2006; Dubrulle and Pourquié, 2004). In contrast, anterior PSM cells are exposed to RA, synthesized by Raldh2 and degraded by Cyp26A1, which promotes expression of adhesion molecules including cadherins and integrins, thereby counteracting Fgf and Wnt signaling (Aulehla and Pourquié, 2010; Duband et al., 1987; Sakai et al., 2001). This shift reduces motility, slows oscillations, and enhances epithelialization and differentiation (Horikawa et al., 1999; Linask et al., 1998; Niederreither and Dollé, 2008). Downstream of Fgf, pathways such as MAPK/Erk and PI3K/Akt regulate motility-associated genes, notably Snail, whose expression peaks in the posterior PSM (Ciruna and Rossant, 2001; Dale et al., 2006; Wahl et al., 2007). At the cellular level, motility is driven by actin cytoskeleton remodeling, tightly controlled by small Rho GTPases (Heasman and Ridley, 2008; Montell, 2008).

Rho family GTPases are central regulators of cytoskeletal dynamics and thereby influence the mechanical properties of cells (Montell, 2008). Through these effects, they control essential behaviors such as cell–cell and cell–matrix adhesion, membrane protrusion formation, and membrane tension regulation (Heasman and Ridley, 2008). Consequently, Rho GTPases affect proliferation, motility, and polarity (Chimini and Chavrier, 2000; Etienne-Manneville and Hall, 2002; Raftopoulou and Hall, 2004). The small Rho GTPases—Rac1, Cdc42, and RhoA— function as molecular switches, cycling between an inactive GDP-bound and an active GTP-bound state. This cycling is controlled by GDP-dissociation inhibitors (GDIs), guanine nucleotide exchange factors (GEFs), and GTPase-activating proteins (GAPs) (Bustelo et al., 2007; Schmidt and Hall, 2002). Once activated, Rho GTPases trigger downstream pathways including PAK, JNK, PI3K, and Par6–aPKC(Aspenström et al., 2004; Nelson, 2003; Ohno, 2001). Among them, Rac1 plays a particularly critical role in early embryonic development. Rac1-null embryos are non-viable (Sugihara et al., 1998), as loss of Rac1 disrupts morphogenetic processes such as neural crest migration and epithelial-to-mesenchymal transition (EMT) during gastrulation. Rac1 deficiency impairs mesodermal migration, disrupts cell polarity, and prevents proper formation of structures such as the anterior visceral endoderm (AVE) and nascent mesoderm (Migeotte et al., 2011; Migeotte et al., 2010; Nakaya et al., 2004).

During paraxial mesoderm development and somitogenesis, Rho GTPases are key regulators of tissue morphogenesis. Rac1 is required for mesodermal cell migration through WAVE-mediated actin remodeling and for somite epithelialization through interactions with factors such as Paraxis, which promote somite boundary formation (Nakaya et al., 2004). All together, these activities ensure accurate segmentation of the paraxial mesoderm and balance between epithelial and mesenchymal states. Cdc42 and RhoA also contribute critically to convergence– extension movements and PSM elongation by modulating adhesion, polarity, and cytoskeletal dynamics (Choi and Han, 2002; Liu and Jessell, 1998). Collectively, Rho GTPases link upstream signals to cellular behaviors that drive segmentation. Notably, these same morphogenetic mechanisms are often co-opted in pathological contexts such as cancer metastasis, underscoring the broader relevance of Rho GTPase function in physiology and disease (Casado-Medrano et al., 2018).

Despite major advances in dissecting the segmentation clock, many molecular regulators of this complex system remain unknown. Even for the well-studied Notch, Wnt, and Fgf pathways, the mechanisms by which they integrate intra- and intercellular cues to drive somitogenesis are not fully resolved. Moreover, the contribution of cellular motility to oscillation dynamics remains poorly understood. Given the established role of Rho GTPases in controlling motility, could they also influence oscillation periodicity? Here, we set out to identify new regulators of the segmentation clock by chemically perturbing signaling pathways involved in motility, polarity, directionality, and mechanosensitivity. Our compound selection was guided both by longstanding questions about motility gradients in the PSM and by emerging evidence linking mechanosensitive pathways to somitogenesis (Bénazéraf et al., 2010; Hubaud et al., 2017). Using a two-dimensional explant culture model derived from the posterior mouse PSM (Lauschke et al., 2013), we show that Rac1 inhibition decreases cell motility and prolongs oscillation periodicity, ultimately disrupting somite formation. Modulation of segmentation clock timing motility can be uncoupled from motility and seems to impact expression of genes involved in Wnt and Notch pathways. These findings uncover a new role for Rac1 activity in regulating segmentation dynamics and highlight broader principles of pathway crosstalk and cell–cell communication in the PSM, with potential implications for other developmental and homeostatic processes.

## Results

### A chemical screen for regulators of segmentation clock periodicity

In mice, the segmentation clock driving somitogenesis oscillates with a periodicity of ∼2 hours. To identify modulators of this timing, we performed a small-scale chemical screen (Richter et al., 2017). Motivated by longstanding questions about the role of the cell motility gradient in the presomitic mesoderm (PSM) (Bénazéraf et al., 2010; Hubaud et al., 2017), we selected compounds targeting cellular processes involved in mechanotransduction, polarity, and directional motility (Supplementary Table 1). Compounds were applied to two-dimensional explant cultures derived from the posterior tip of the mouse PSM (Lauschke et al., 2013)(Figure 1A). Segmentation clock oscillations were monitored using the LuVeLu (LVL) fluorescent reporter, which reflects periodic *Lfng* expression in PSM cells (Aulehla et al., 2008). This primary culture system recapitulates PSM dynamics and provides a robust platform for chemical screening. Time-lapse imaging was followed by quantitative analysis of collective fluorescence dynamics using a standardized pipeline (Figure 1A, see Methods).

**Figure 1.**
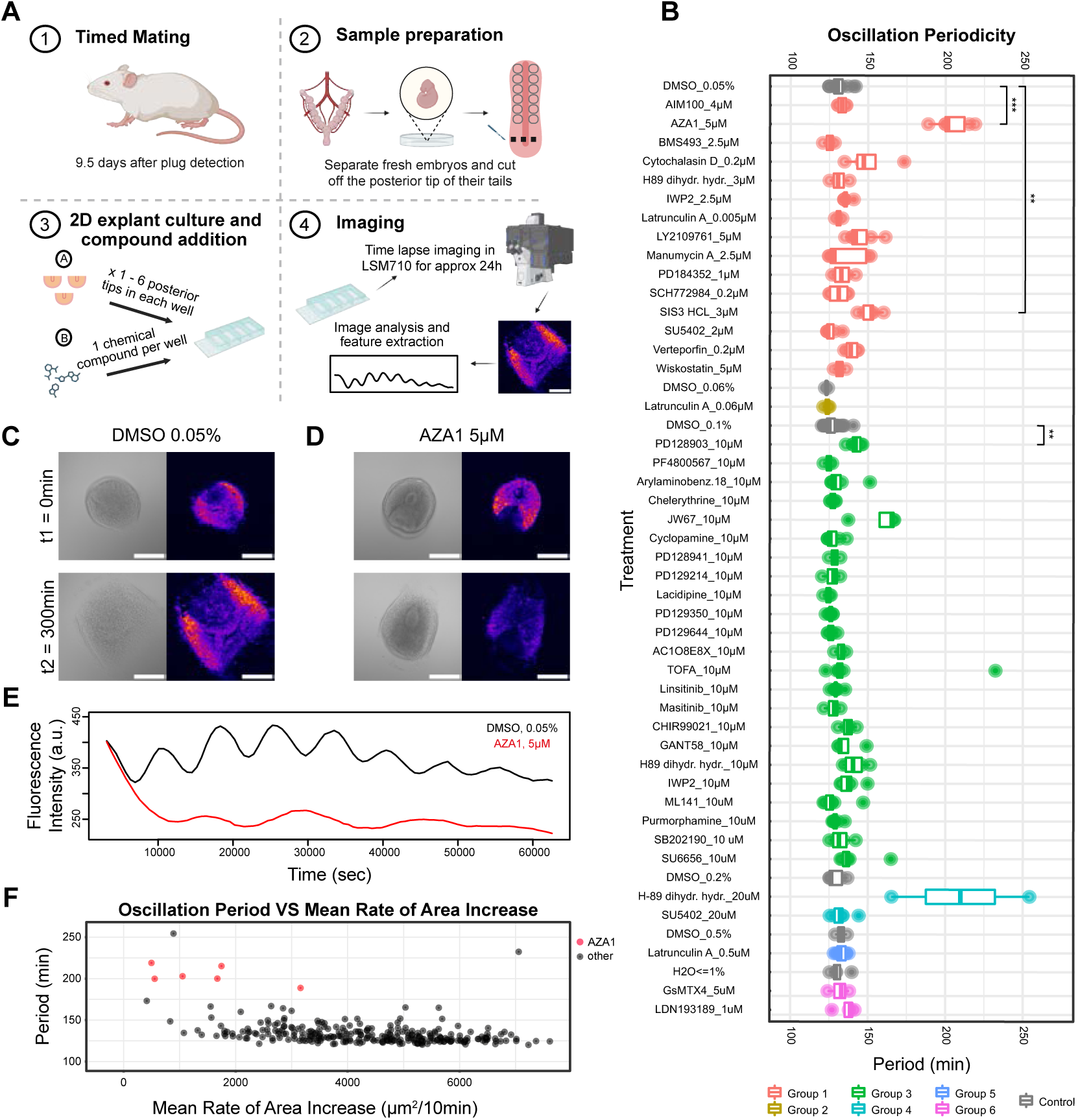
A small molecule screen identifies AZA1 (5 μM) as a potent modulator of oscillation period and tissue spreading in mPSM explants. **(A)** Schematic representation of the chemical screen experimental setup using a 2D explant culture assay (see Methods). Figure created with BioRender. **(B)** Summary of periodicity measurements from the small molecule screen. Compounds are grouped by vehicle control type and concentration (see Methods; compound details in Table 1). Controls are shown in grey. **(C)** Representative images of mouse presomitic mesoderm (mPSM) explants cultured on fibronectin-coated plates and treated with vehicle control (0.05% DMSO). Brightfield (left) and YFP fluorescence images (right) from LuVeLu reporter explants at the start of imaging (t₁ = 0 min) and after 5 hours (t₂ = 300 min). Over time, explants spread and display outward-traveling waves of LuVeLu expression (see Supplementary Video 1). **(D)** Representative images of mPSM explants treated with 5 μM AZA1 under identical conditions. AZA1 does not impair attachment to the fibronectin coated glass or cause overt toxicity (see Supplementary Video 2). **(E)** Quantification of LuVeLu fluorescence intensity over time in DMSO (black) and AZA1-treated (red) explants. Oscillatory dynamics are maintained under both conditions, but AZA1 treatment results in a marked increase in the oscillation period. **(F)** Scatterplot of oscillation period versus mean explant spreading rate for all compounds tested. Each dot represents one mPSM explant. AZA1-treated samples are shown in red. A direct correlation between period and spreading is not observed. All images in panels (C–D) were processed with identical brightness and contrast settings. Scale bars: 200 μm.

To estimate oscillation period, we applied Singular Spectrum Analysis (SSA) (Golyandina and Korobeynikov, 2014; Zhigljavsky and Golyandina, 2013) (Figure S1). Unlike traditional methods, SSA does not require prior knowledge of the expected period range, making it particularly well suited for unbiased screening. SSA also decomposes the signal to quantify the contribution of individual components to the overall time series. In parallel, we measured explant area expansion as a proxy for tissue mechanical properties, primarily reflecting cell migration across the fibronectin-coated substrate (Figure S2). For each treatment, effects on oscillation periodicity and motility were compared with matched controls, applying False Discovery Rate (FDR) correction for multiple testing (Benjamini and Hochberg, 1995).

Control data from independent experiments were pooled for each concentration group and compared against corresponding compound treatments (Figure 1B). As expected, DMSO treatment did not alter oscillation period relative to previously reported values (Lauschke et al., 2013). The most striking hit was Aza1 (Figure 1B), a dual Rac1/Cdc42 inhibitor (Zins et al., 2013). Treatment with 5 µM Aza1 significantly lengthened the LVL oscillation period, from ∼140 minutes in controls to 180–230 minutes (Figure 1C–E, Suppl. Videos 1, 2). Consistent with Rac1/Cdc42 inhibition, Aza1 also strongly reduced tissue spreading, indicating impaired motility (Figure 1C–D,F; Figure S3; Suppl. Videos 1, 2). Several other compounds also reduced spreading, while two—Wiskostatin and BMS493—unexpectedly increased it (Figure S3). Notably, changes in motility did not consistently correlate with oscillation period. For example, cytochalasin almost completely blocked spreading yet had no detectable effect on LVL periodicity (Figure 1B, Figure S3). Plotting oscillation period against spreading rate across all treatments revealed no clear relationship (Figure 1F), suggesting that changes in motility alone cannot explain alterations in segmentation clock timing.

### Rac1 inhibition increases oscillation period

Because Aza1 inhibits both Rac1 and Cdc42, we next asked which GTPase is primarily responsible for the observed increase in oscillation periodicity. To distinguish their roles, we tested compounds targeting either Rac1, Cdc42, or both. The selective Cdc42 inhibitors ML141 and CASIN (Liu et al., 2019; Surviladze et al., 2010) produced only minor effects. At low concentration (20 µM), ML141 caused a modest but statistically significant period increase, but this effect disappeared at higher doses and was not reproduced with CASIN (Figure 2A). These results suggest that Cdc42 contributes little, if at all, to segmentation clock periodicity and cannot account for the strong Aza1 phenotype.

**Figure 2.**
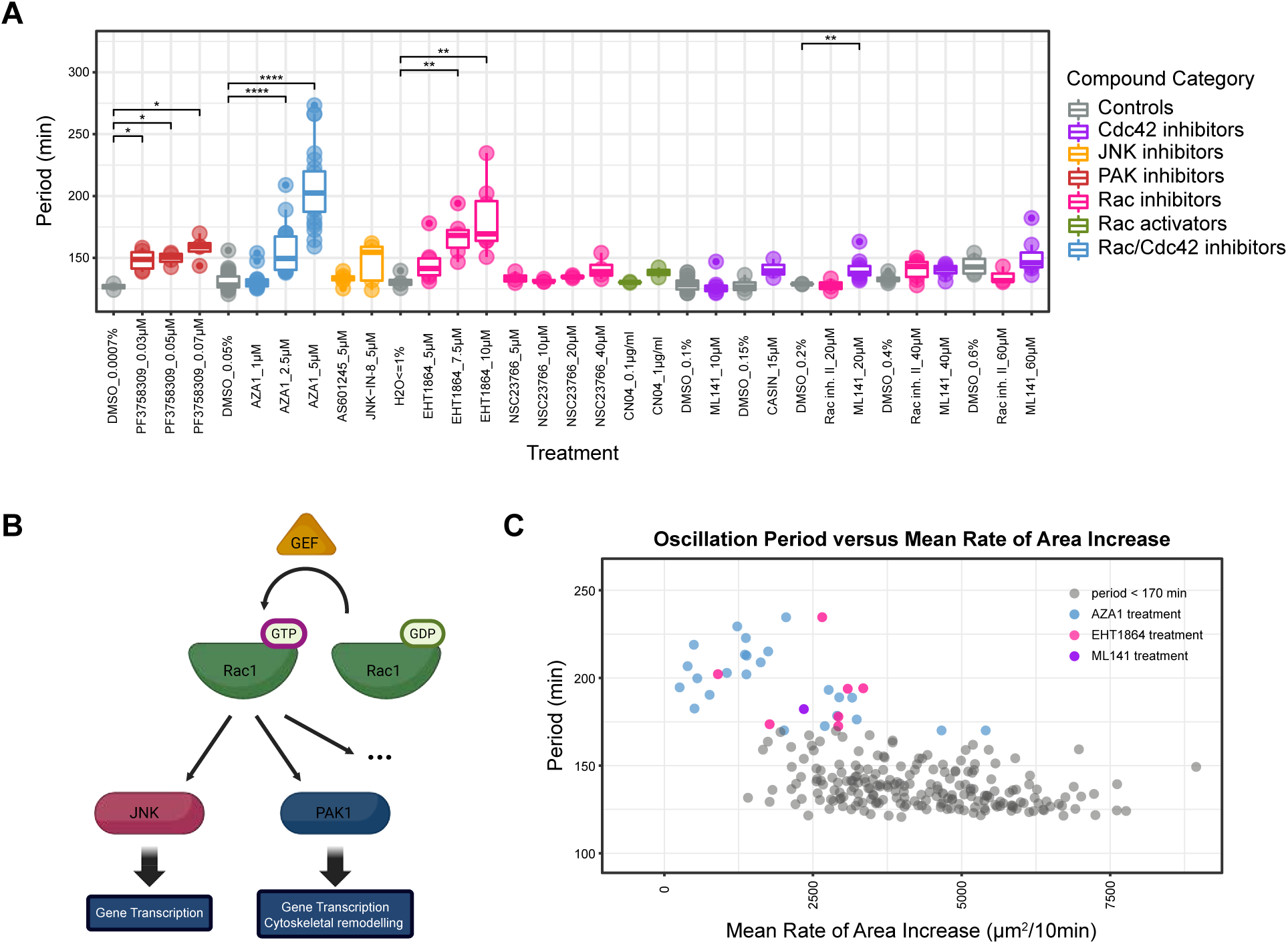
Oscillation dynamics and tissue spreading are differentially regulated by Rac1/Cdc42 pathway modulators. **(A)** Oscillation periodicity measurements in mPSM explants treated with compounds targeting the Rac1 and Cdc42 small GTPase signaling pathways (compound details in Table 1). Color coding reflects the compound’s mode of action: controls (grey), Rac1/Cdc42 dual inhibitors (light blue), Cdc42-selective inhibitors (purple), Rac1-selective inhibitors (pink), Rac1 activators (green), PAK inhibitors (red), and JNK inhibitors (orange). **(B)** Simplified schematic of Rac1 being activated by GEFs through GDP-GTP exchange. Once activated, Rac1 signals, among other pathways, via JNK to activate gene transcription and/or via PAK1 to additionally regulate cytoskeletal remodeling. **(C)** Scatterplot of oscillation period versus mean rate of area increase in explants treated with Rac1/Cdc42-associated compounds. Each point represents a single mPSM sample. Colored dots indicate samples with oscillation periods exceeding 170 minutes: pink (EHT1864), light blue (AZA1), and purple (ML141). AZA1 and EHT1864 both lead to increased oscillation periodicity and reduced spreading.

In contrast, the selective Rac1 inhibitor EHT1864 (Shutes et al., 2007) robustly recapitulated the Aza1-induced period lengthening. Both Aza1 and EHT1864 showed dose-dependent effects, with higher concentrations producing greater extensions (Figure 2A). Interestingly, other Rac1 inhibitors—including NSC23766, the parent compound of Aza1 (Gao et al., 2004; Zins et al., 2013)—did not significantly affect oscillations. Likewise, Rac1 activation by CN04 (Lerm et al., 1999) did not shorten the period below control levels, suggesting an intrinsic lower limit to how fast oscillations can occur. Together, these findings highlight Rac1 as a specific and critical regulator of segmentation clock periodicity.

Differences among Rac1-targeting compounds likely reflect their distinct mechanisms of action. NSC23766 and Rac1 inhibitor II, which had no effect on periodicity, block Rac1 activation by preventing interaction with guanine nucleotide exchange factors (GEFs) such as TIAM1. By contrast, EHT1864 interferes with Rac1’s ability to activate downstream effectors, particularly p21-activated kinases (PAKs) (Shutes et al., 2007). This mechanistic divergence may explain why only some Rac1 inhibitors extend periodicity, underscoring the importance of downstream signaling context in PSM cells (Figure 2B). Consistent with this model, direct inhibition of the Rac1–PAK axis using PF3758309 significantly increased oscillation period at all concentrations tested (Figure 2A). Conversely, inhibition of c-Jun N-terminal kinases (JNKs), another Rac1 effector branch, with JNK-IN-8 and AS601245 had no effect. These data indicate that Rac1 modulates segmentation clock timing primarily via PAK signaling.

Since Rac1 and Cdc42 both regulate cytoskeletal dynamics and motility, we also assessed their effects on PSM spreading. High doses of Aza1 or ML141 markedly reduced explant expansion, consistent with impaired motility (Figure S4). Notably, only Rac1 inhibition (Aza1 or EHT1864) lengthened oscillation period, reinforcing Rac1’s specific role in clock regulation. Furthermore, PAK inhibition extended periodicity without reducing spreading, while Aza1 and EHT1864 affected both (Figure 2C). Thus, motility changes alone cannot fully account for altered periodicity. We conclude that Rac1 regulates oscillation timing at least in part through PAK signaling, with additional mechanisms—possibly involving motility—also contributing.

Because our measurements capture population-level dynamics, we considered whether the apparent period increase might result from loss of synchrony between PSM cells. Reduced synchrony could appear as slower oscillations at the tissue scale (Roth et al., 2023; Winfree, 1980). To test this, we used an alternative assay in which PSM tissue is dissociated, reaggregated, and allowed to self-organize into emergent PSM (ePSM) cultures (Tsiairis and Aulehla, 2016) (Figure 3A). These cultures reproduce key features of native PSM dynamics, including synchronized oscillations and traveling waves. We compared controls with ePSMs treated with 2.5 µM or 5 µM Aza1 (Figure 3B,C). Given Rac1/Cdc42’s roles in motility, we expected impaired sorting and disrupted synchrony. Surprisingly, dual inhibition did not block self-organization or wave formation. Oscillatory foci still emerged but tended to cease oscillations prematurely compared with controls (Figure 3B; Suppl. Videos 3–5).

**Figure 3.**
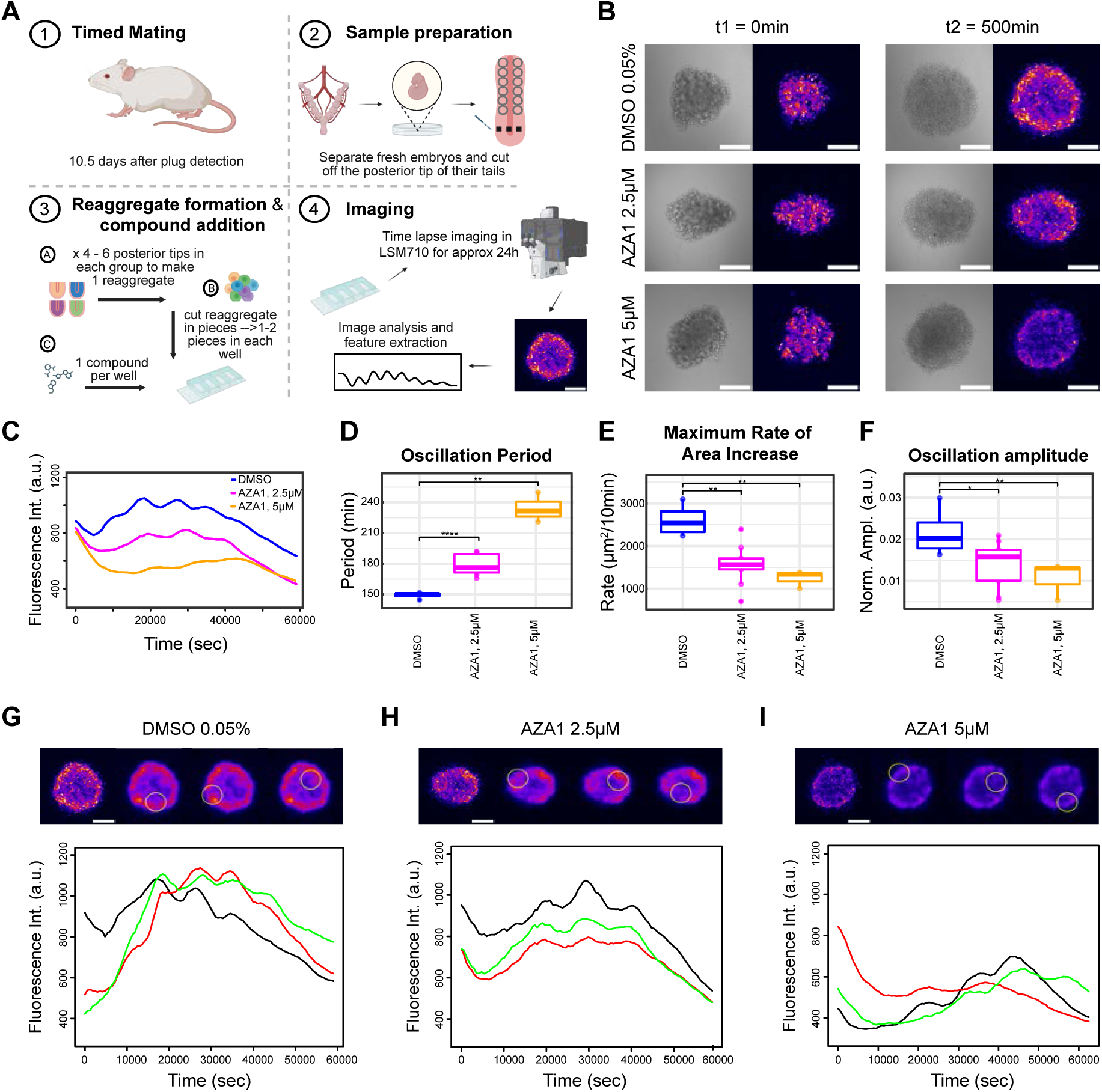
AZA1 treatment alters the temporal and spatial dynamics of ePSMs but does not disrupt oscillatory synchrony. **(A)** Schematic overview of the reaggregation assay used to generate engineered presomitic mesoderms (ePSMs) from E10.5 mouse embryos, followed by compound addition and time-lapse imaging (see Methods). Illustration created with BioRender. **(B)** Representative brightfield (left) and YFP-channel (right) images of ePSMs treated with DMSO (0.05%), AZA1 (2.5 μM), or AZA1 (5 μM), shown at the start of imaging (t₁ = 0 min) and after 500 min (t₂ ≈ 8 h 20 min) (see Supplementary Videos 3-5). **(C)** Mean fluorescence intensity over time for the ePSMs shown in (B), indicating altered dynamics in AZA1-treated conditions. **(D–F)** Quantifications of oscillatory parameters reveal a concentration-dependent effect of AZA1: (D) increase in oscillation period, (E) decrease in maximum rate of area increase, and (F) reduction in oscillation amplitude. **(G–I)** Spatial synchrony analysis of local oscillatory activity within the ePSMs shown in (B). YFP images at t₂ = 500 min display selected circular regions of interest (yellow circles), with corresponding fluorescence intensity traces plotted below. Despite changes in temporal dynamics, oscillations remain synchronized across spatial domains in all treatment conditions. A Gaussian filter (radius = 10 μm) was applied for quantification (see Methods). All images were processed using identical brightness and contrast thresholds to enable direct comparison. Scale bars: 200 μm.

Global analysis revealed dose-dependent decreases in reporter intensity and progressive lengthening of oscillation period under Aza1 treatment (Figure 3D). As previously reported, ePSM periods were longer than explants under control conditions(Tsiairis and Aulehla, 2016). Similar to explants, Aza1 slowed spreading of ePSMs on fibronectin-coated glass (Figure 3E) and reduced oscillation amplitude (Figure 3F). To test whether amplitude loss reflected desynchronization, we examined subregions of individual ePSMs. Control cultures showed synchronized peaks and troughs across regions (Figure 3G), and synchrony was maintained under Aza1 treatment despite increased period and reduced amplitude (Figure 3H,I). Thus, Rac1/Cdc42 inhibition does not impair synchrony or wave propagation. Instead, Aza1 modifies oscillation dynamics while preserving cellular self-organization.

### Phase resetting due to Rac1 inhibition

Given that Aza1 treatment consistently produced up to a twofold increase in oscillation period across both mPSM and ePSM cultures, we next asked whether this effect is reversible upon compound removal. Specifically, we tested whether Rac1 inhibition induces only a transient shift in oscillator dynamics or a more persistent reprogramming of the system. To address this, we performed a washout experiment in which cultures were treated with either DMSO or Aza1 for six hours, followed by medium replacement with fresh, compound-free medium. Imaging was then continued for ten additional hours, enabling continuous monitoring of individual samples before and after washout alongside time-matched controls (Figure 4A–B, Suppl. Videos 6–8). Following removal of Aza1, oscillations accelerated, with periodicity returning to near-control values (Figure 4C–E). This recovery was most pronounced after removal of 5 µM Aza1, where periods decreased substantially relative to pre-wash values and closely matched DMSO controls subjected to the same protocol (Figure 4F). Thus, Rac1 inhibition alters segmentation clock timing in a reversible manner and does not permanently reprogram oscillator behavior.

**Figure 4.**
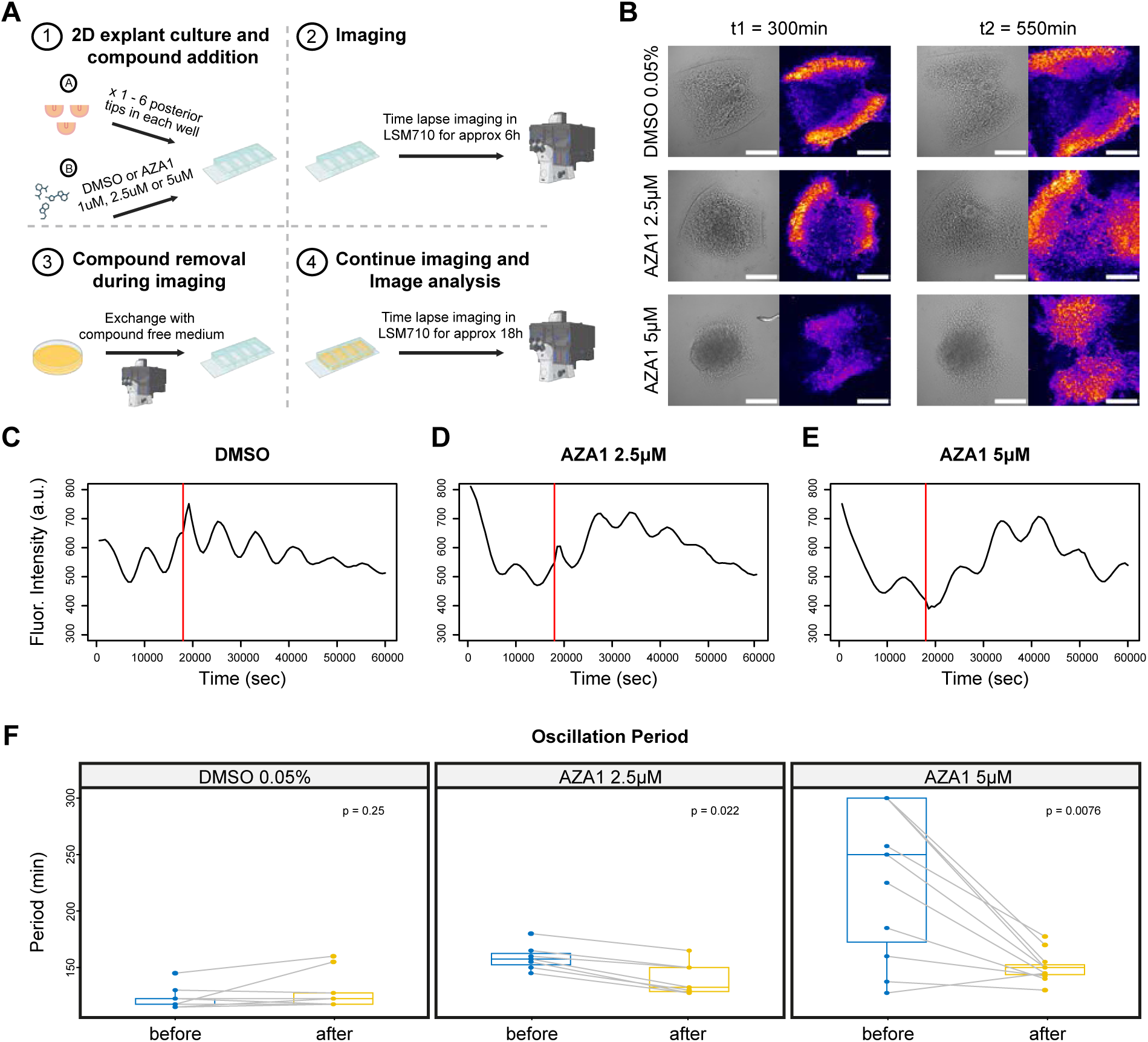
The oscillation period in mPSMs recovers after AZA1 washout. **(A)** Schematic representation of the experimental setup for AZA1 treatment and subsequent removal in the mPSM primary culture assay (see Methods). Illustration created with BioRender. **(B)** Representative brightfield (left) and YFP-channel (right) images of mPSMs treated with DMSO (0.05%), AZA1 (2.5 μM), or AZA1 (5 μM), shown immediately before compound removal at t₁ = 300 min (5 h) and some time after washout at t₂ = 550 min (≈9 h 10 min). Following removal, a recovery of fluorescence intensity is observed in AZA1-treated samples (see Supplementary Videos 6-8). **(C–E)** Plots of mean fluorescence intensity over time for the DMSO-, 2.5 μM AZA1-, and 5 μM AZA1-treated mPSMs shown in (B). The time of compound removal is indicated by a red line. A transient spike immediately following removal corresponds to a minor z-plane shift during imaging and is considered an artefact introduced by the manipulation. **(F)** Quantification of oscillation periods before and after compound removal. Washout of AZA1 results in a partial restoration of the oscillation period. All images were processed with consistent brightness and contrast settings for accurate comparison. Scale bars: 200 μm.

Closer inspection of oscillatory trajectories revealed a smooth continuity of phase progression across the washout, suggesting that phase transitions remained intact (Figure S5). This observation prompted us to investigate how Rac1 inhibition influences oscillator phase. We asked whether introducing Aza1 during ongoing oscillations perturbs phase in a specific, non-random manner. To test this, we added Aza1 to untreated mPSM cultures during live imaging, six hours after imaging onset, and tracked the dynamics (Figure 5A–B, Suppl. Videos 9–11). Addition of 5 µM Aza1 triggered a rapid rise in LuVeLu signal amplitude, followed by the establishment of a new steady state with low-amplitude, long-period oscillations—consistent with continuous Aza1 exposure from culture onset (Figure 1E, Figure 5E–F, S6). A milder but measurable period increase was observed with 2.5 µM Aza1, whereas DMSO-treated controls exhibited only a gradual, modest period extension (Figure 5C–F). This slow extension in controls likely reflects the intrinsic maturation of PSM tissue, as cells progressively acquire anterior-like identities associated with slower oscillations (Tsiairis and Aulehla, 2016).

**Figure 5.**
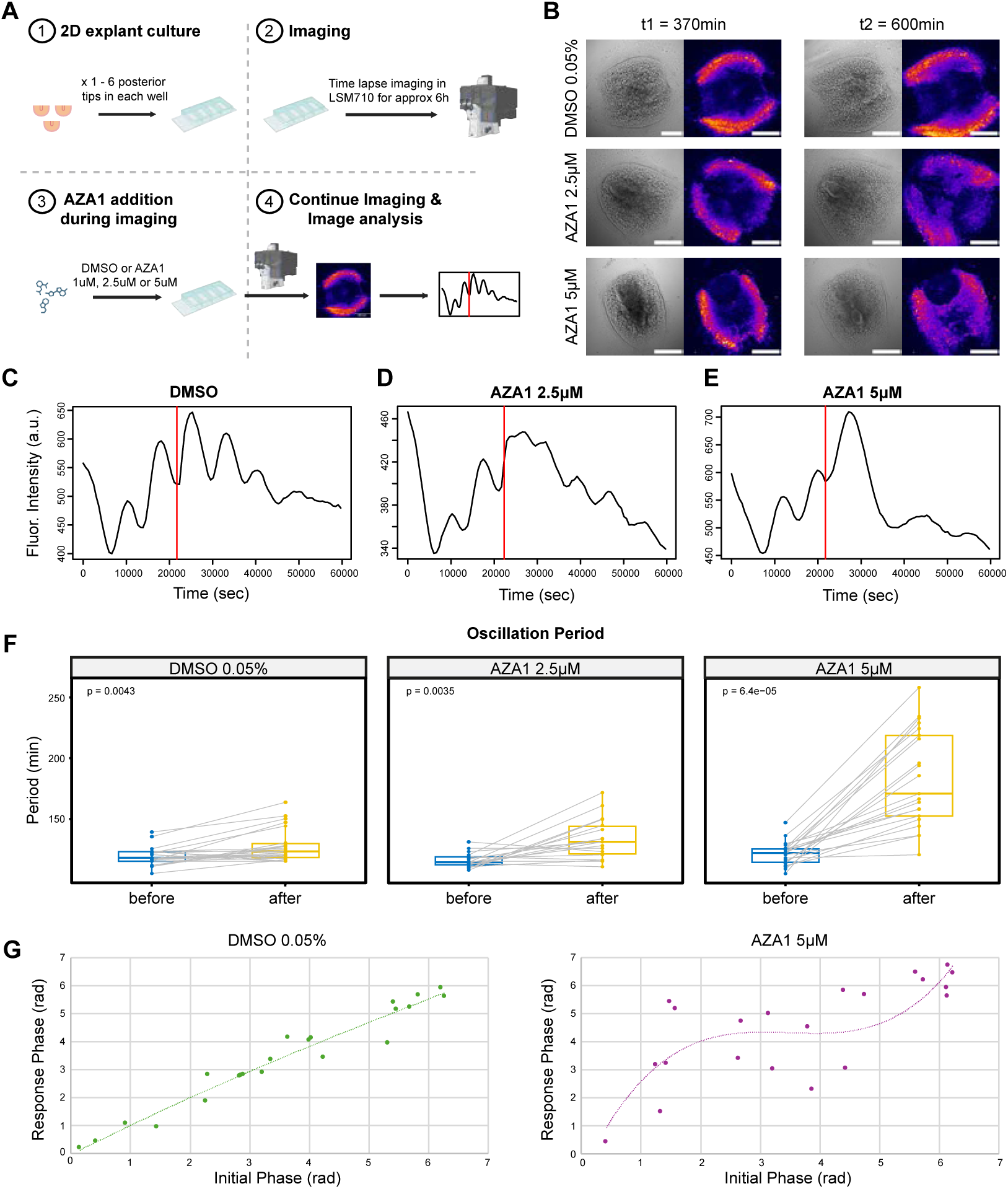
AZA1 treatment induces an immediate reset of oscillation phase and period. **(A)** Schematic overview of the experimental setup for AZA1 treatment late addition in the mPSM primary culture assay (see Methods). Illustration created with BioRender. **(B)** Representative images of mPSM explants treated with DMSO (control), 2.5 μM AZA1, or 5 μM AZA1. Left panels show brightfield images; right panels show YFP fluorescence. Images were captured immediately before compound addition (t₁ = 370 min, 6 h 10 min, after imaging start) and at t₂ = 600 min (10 h after imaging start) (see Supplementary Videos 9-11). **(C–E)** Time course plots of mean fluorescence intensity for DMSO-, 2.5 μM AZA1-, and 5 μM AZA1-treated samples corresponding to the conditions in (B). The time of compound addition is marked by a red vertical line. Notably, 5 μM AZA1 triggers an immediate reset of the oscillation phase accompanied by a transient increase in fluorescence intensity. **(F)** Quantification of oscillation periods before and after treatment with DMSO, 2.5 μM AZA1, or 5 μM AZA1 in mPSMs. **(G)** Phase Resetting Curves measuring the impact of adding DMSO or 5µM AZA1 on the oscillatory dynamics of mPSM cultures. Scale bar: 200 μm.

To characterize the immediate oscillator response to Aza1, we analyzed the relationship between oscillation phase before and after treatment, generating phase response curves (PRCs)—a classical approach to quantify oscillator resetting (Johnson, 1999; Winfree, 1980). In DMSO-treated controls, pre- and post-treatment phases remained aligned, producing a near-linear identity relationship (Figure 5E). In contrast, 5 µM Aza1 caused a significant deviation from this diagonal, consistent with a phase reset. The winding number remained one, indicating that a complete oscillatory cycle occurred, but the phase skewed preferentially toward values greater than π (Figure 5E)—corresponding to the late portion of the cycle, between baseline and peak. This reveals that Aza1 induces a strong, directed phase reset confined to a specific window of the cycle.

In summary, Rac1 inhibition by Aza1 not only lengthens the segmentation clock period but also actively resets oscillator phase, thereby altering the trajectory of PSM dynamics in phase space. These findings suggest that Rac1-dependent pathways contribute to phase entrainment of the segmentation clock, adding a new layer of temporal regulation beyond simple modulation of periodicity.

### Rac1 inhibition impact on gene expression

Oscillatory gene expression in the presomitic mesoderm (PSM) is coordinated by Notch, Wnt, and FGF signaling pathways. To examine how Rac1 inhibition affects these pathways, we used an in vitro culture system that supports growth and segmentation of PSM explants (Lauschke et al., 2013). Posterior embryo fragments—including two pre-formed somites—were dissected and cultured for 21.5 hours in the presence of 5 µM Aza1 or DMSO control (Figure 6A). At the end of the culture period, newly formed somites along with the first two pre-existing ones were separated from the PSM, and gene expression in the remaining PSM tissue was analyzed by RT-PCR. This analysis revealed pathway-specific effects: *Hes5*, a Notch target, was upregulated, whereas *Msgn1*, a Wnt effector, was downregulated in Aza1-treated samples (Figure 6B, Figure S7). In contrast, *Dusp4*, an FGF target, remained unchanged. These results indicate that Rac1 inhibition selectively perturbs Notch and Wnt signaling, potentially through direct pathway modulation or via known crosstalk between these pathways (Goldbeter and Pourquié, 2008).

**Figure 6.**
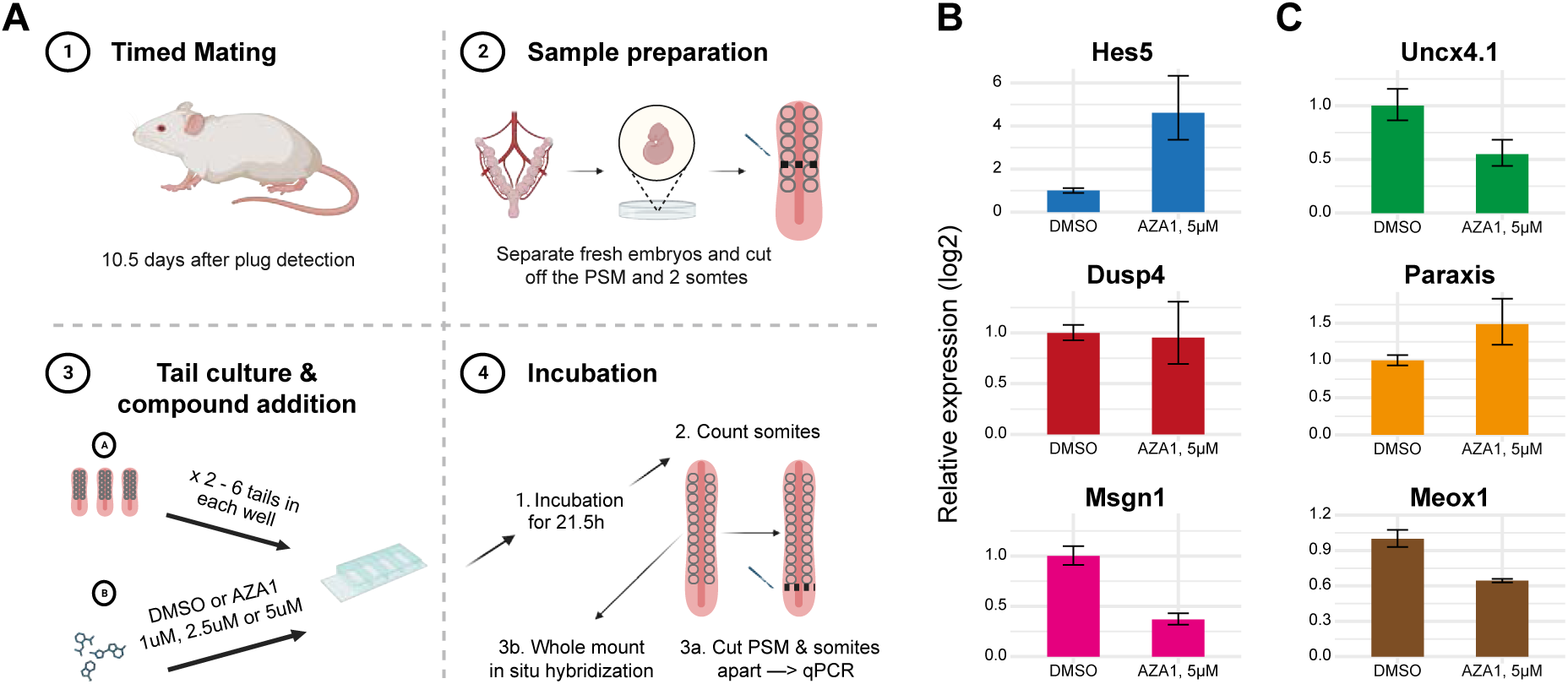
Rac1 inhibition induces transcriptional changes in both somitic and PSM tissues. **(A)** Schematic of the experimental setup for tail explant experiments (see Methods), created with BioRender. **(B)** Quantification of relative expression levels of PSM markers (*Hes5*, *Dusp4*, and *Msgn1*) by qPCR in tail explants treated with DMSO or 5 μM AZA1. **(C)** Quantification of relative expression levels of somitic markers (*Uncx4.1*, *Paraxis*, and *Meox1*) by qPCR under the same conditions.

Given the impact on oscillatory dynamics, we next assessed how Rac1 inhibition affects somite formation and patterning. Using RT-PCR, we measured expression of key somitic markers: *Paraxis*, broadly expressed in somites and required for epithelialization (Rowton et al., 2013); *Uncx4.1*, a posterior somite marker (Mansouri et al., 2000); and *Meox1*, marking the dorsomedial domain (Reijntjes et al., 2007; Skuntz et al., 2009). Among these, *Paraxis* expression was elevated, albeit variably across replicates, suggesting enhanced or prolonged epithelialization (Figure 6C, Figure S7). In contrast, both *Uncx4.1* and *Meox1* were downregulated, indicating defects in anterior–posterior and dorsoventral somite patterning (Figure 6C, Figure S7). Although somites still formed under Aza1 treatment, these molecular changes suggest disrupted spatial organization of somite compartments, which could compromise the subsequent differentiation of the dermomyotome, sclerotome, and other derivatives.

### Somite formation defects after increased periodicity

To directly assess morphogenetic outcomes, we used the in vitro PSM culture system to monitor somite formation during Rac1 inhibition (Figure 6A). Explants were cultured over several hours, allowing new somites to form sequentially posterior to the pre-existing pairs (Figure 7A,B). Somite boundaries were visualized via in situ hybridization for *Uncx4.1*, which marks the posterior half of each somite (Figure S8). We proceeded to quantify the number of somites in each experimental group.

**Figure 7.**
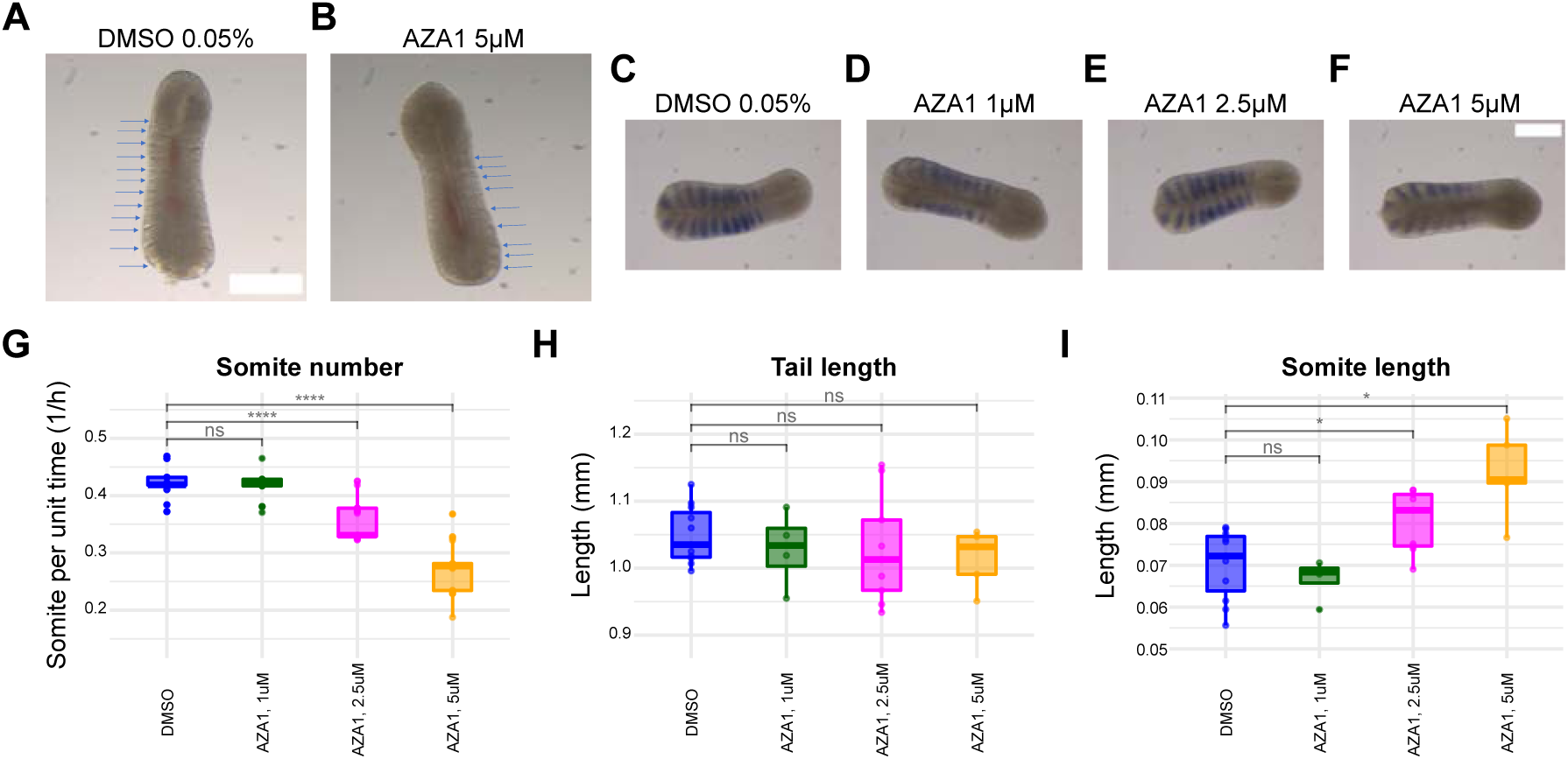
Rac1 inhibition alters somitogenesis in tail explants. **(A–B)** Tail explants cultured for 21.5 hours with DMSO (A) or 5 μM AZA1 (B) and imaged thereafter. Arrows indicate somites. AZA1-treated explants show fewer somites with visibly distorted morphology and poorly defined boundaries. **(C–F)** In situ hybridization for *Uncx4.1* in tail explants cultured for 21.5 hours with 0.05% DMSO (C) or increasing concentrations of AZA1: 1 μM (D), 2.5 μM (E), and 5 μM (F). **(G–I)** Quantification of (G) somite number, (H) total tail length, and (I) average somite length across treatments. Rac1 inhibition leads to a concentration-dependent increase in somite length without affecting total tail length, resulting in fewer somites. Statistical analyses comparing treatment groups are described in Methods. Scale bar: 300 μm.

Aza1 treatment, and thus Rac1 inhibition, caused a clear, dose-dependent reduction in the number of somites formed during the culture period (Figure 7C–G). Two mechanistic scenarios could explain this outcome: (1) a slower growth rate of the explant, which, in combination with the slower segmentation clock, produces fewer, normally sized somites and a shorter PSM, or (2) a normal axial growth rate, which together with the slower segmentation clock, generates fewer but larger somites, preserving total PSM length. To distinguish these possibilities, we measured both overall explant length (Figure 7H) and individual somite length (Figure 7I). Results supported the second scenario: somites formed under Aza1 treatment were significantly larger, while total explant length remained comparable to controls.

These findings align with predictions of the clock-and-wavefront model of somitogenesis, which posits that somite size is determined by the oscillation period relative to wavefront progression. Accordingly, lengthening the oscillation period—as observed with Rac1 inhibition—produces fewer, larger somites without affecting overall axial growth. Collectively, these results demonstrate that Rac1 perturbation disrupts both molecular patterning and morphological segmentation, highlighting its critical role in coordinating the tempo and fidelity of somite formation.

## Discussion

Our study identifies Rac1 as a novel regulator of segmentation clock period in the mouse presomitic mesoderm (PSM). Using a chemical screen, we found that Rac1 inhibition nearly doubled oscillation period in 2D posterior PSM explants—an unprecedented effect in this system. Transient treatment with AZA1, a Rac1/Cdc42 inhibitor, shifted the system to a new steady state characterized by high-period, low-amplitude oscillations, while subsequent recovery upon AZA1 washout argues against major toxicity effects. This regulation likely occurs through the Rac1 effector PAK (Aspenström et al., 2004), as PAK inhibition also extended the period, though to a lesser degree. Consistent with these changes, AZA1-treated tail explants formed fewer somites with irregular boundaries.

Rho-family GTPases are well-established regulators of development and disease (Boettner and Van Aelst, 2002; Gomez del Pulgar et al., 2005; Rossman et al., 2005; Sahai and Marshall, 2002; Vega and Ridley, 2008). Biochemical studies indicate that Cdc42 and Rac1 act through either shared or distinct pathways depending on cellular context (Fukata et al., 2003; Schock and Perrimon, 2002; Settleman, 2001; Van Aelst and Symons, 2002). In our experiments, selective Cdc42 inhibition did not significantly alter oscillation period, whereas selective Rac1 inhibition with EHT1864 or dual Rac1/Cdc42 inhibition with AZA1 markedly prolonged it (Figures 1, 2). These findings suggest a predominant role for Rac1 in regulating the segmentation clock. We focus on Rac1 in the subsequent discussion, while noting that dual inhibition may contribute to the somitic defects observed with AZA1. The differential effects of Cdc42 and Rac1 provide new insight into the specific roles of these GTPases in morphogenetic processes during early embryogenesis.

A key challenge in dissecting Rho GTPase function in the segmentation clock is identifying which of the ∼80 characterized GEFs are specifically expressed in mouse PSM. GEFs mediate GDP–GTP exchange, activating small GTPases (Gao et al., 2004). The Rac1 inhibitors in our screen act via distinct mechanisms, targeting either specific GEF interactions or downstream effectors. Rac1 Inhibitor II and NSC23766 block Rac1–Tiam1 interaction without affecting Cdc42, RhoA, PAK1 activation, or Rac1 interactions with other GEFs such as Vav or Lbc (Blangy et al., 2000; Debant et al., 1996; Gao et al., 2004). In contrast, EHT1864 blocks GTP binding to all Rac family members (Rac1, Rac1b, Rac2, Rac3), inhibiting interactions with all effectors including PAK1 (Shutes et al., 2007). AZA1, a dual Rac1/Cdc42 inhibitor, reduces phosphorylation of PAK1/2, Akt, and BAD but not ERK, JNK, or p38 (Zins et al., 2013). Selective disruption of Rac1–Tiam1 interaction (Rac1 Inhibitor II, NSC23766) does not alter oscillation period, whereas broader inhibition (EHT1864, AZA1) significantly prolongs it.

These pharmacological distinctions raise the question of how Rac1 inhibition mechanistically alters segmentation clock dynamics. One possibility is an effect on cell cycle dynamics, as Rac1 and Cdc42 regulate mitosis (Hirata et al., 2020; Michaelson et al., 2008; Walmod et al., 2004). Cell division introduces noise into the segmentation clock, with daughter cells resynchronizing post-mitosis (Delaune et al., 2012). Rac1 inhibition could prolong the cell cycle or disrupt post-mitotic synchronization, contributing to period lengthening; comparing cell cycle duration under DMSO and AZA1 would clarify this link.

A second possibility is weakened coupling between PSM cells, promoting desynchronization and slowing ensemble oscillations (Herrgen et al., 2010; Masamizu et al., 2006). However, in our analysis, Aza1-treated cells still generate waves of gene expression in mPSM cultures and coordinate these behaviors in distinct domains in ePSM cultures, arguing against a desynchronization action.

Third, Rac1’s central role in cell mechanics—including motility and adhesion—could indirectly affect oscillatory behavior. Changes in cell shape or migration, as observed in fibronectin-cultured PSM cells (Hubaud et al., 2017), might influence the clock. In our study, both AZA1 and SIS3 reduced the spreading area of 2D mPSM explants and increased oscillation period, whereas other inhibitors showed no consistent correlation between spreading and periodicity (Figures 6F, 8A, 13). The Cdc42 inhibitor ML141 perturbed motility without affecting period, while the PAK inhibitor PF-3758309 affected period but not spreading. This divergence highlights distinct roles for Rac1 and Cdc42 in PSM dynamics. Given context-dependent signaling through shared or separate effectors (Etienne-Manneville and Hall, 2002; Van Aelst and Symons, 2002), our results support a dominant role for Rac1 over Cdc42 in maintaining normal oscillation period and motility. Overall, reduced motility may coincide with increased periodicity in some cases, but these phenomena are not strictly linked.

If motility alone cannot explain the period increase, how does Rac1 regulate oscillation dynamics? Our qPCR analysis of tail explants showed that AZA1 downregulates *Msgn1*, a Wnt target (Wittler et al., 2007; Yamaguchi et al., 1999), consistent with previous reports linking Rac1 to Wnt regulation (Habas et al., 2003). In limb development, Rac1 activates Wnt/β-catenin via JNK2-mediated β-catenin phosphorylation, with Rac1 knockout phenocopying β-catenin deletion (Wu et al., 2008). Given parallels between limb and PSM patterning (Aulehla and Pourquié, 2010; Sheeba et al., 2016; Tabin and Wolpert, 2007), we hypothesize that Rac1 regulates Wnt activity in PSM via PAK1 rather than JNK, as PAK but not JNK inhibitors altered oscillation period. In other systems PAK proteins phosphorylate β-catenin at Ser675 to prevent degradation, promote nuclear localization and TCF/LEF transcriptional activity, and enhance chromatin binding via SETD6-mediated methylation of PAK4 (Levy et al., 2011; O’Neill et al., 2014; Vershinin et al., 2016). It is also known that in the PSM Wnt inhibition increases oscillation period, and Wnt and Notch pathways are co-dependent (Gibb et al., 2009). We hypothesize that, by deactivating Wnt, AZA1 may shift PSM cells toward a more anterior identity, resulting in slower oscillations (Lauschke et al., 2013). These findings support a model in which Rac1–PAK signaling activates Wnt/β-catenin to regulate segmentation clock periodicity.

An open question is why Rac1 inhibition produces a larger period increase than PAK inhibition. Our broad-spectrum PAK inhibitor targeted PAK1–6; future experiments should test selective PAK1–3 inhibition. If Rac1–PAK modulates period via Wnt, Wnt inhibition should also increase periodicity. While IWP-2 showed a trend toward longer periods, this effect was not significant, possibly due to suboptimal concentration or limited sample size. AZA1 also increased Notch target *Hes5*, suggesting broader Wnt and Notch modulation. Combined effects on both pathways, potentially synergized with reduced motility, may explain why AZA1 nearly doubles the oscillation period.

Beyond periodicity, AZA1 rapidly resets oscillator phase, as revealed by phase response curve (PRC) analysis. PRCs quantify phase-dependent effects of perturbations on oscillators (Johnson, 1999). AZA1 induced resetting toward a specific cycle phase, corresponding to increasing LuVeLu signal, producing a PRC with winding number +1. This profile suggests that targeted Rac1 modulation could synchronize or entrain PSM cells.

Although Rac1 is best known for its early role in gastrulation, our findings highlight its influence on oscillations in presomitic mesoderm (PSM) cells and its established importance—alongside Cdc42—in later developmental stages, including somite formation and patterning. In the anterior PSM, Rac1/Cdc42 activity is essential for the mesenchymal-to-epithelial transition (MET) that generates somite epithelial structures (Nakaya et al., 2004; Takahashi et al., 2005). Paraxis, a transcription factor regulating MET, has been proposed to modulate Rac1 activity through GEFs, GAPs, or GDIs, requiring precise control for proper epithelialization (Nakaya et al., 2004). While previous work reported that Rac1 deletion did not alter Paraxis expression, we observed upregulation of Paraxis mRNA in AZA1-treated tail explants, despite the formation of fewer but larger somites. This suggests a feedback mechanism whereby reduced Rac1 activity induces Paraxis expression to restore Rac1 signaling. Given that Wnt/β-catenin can activate Paraxis (Linker et al., 2005), and that Paraxis promotes cytoskeletal effectors downstream of Rac1/Cdc42 (Daggett et al., 2004; Smallhorn et al., 2004)(Daggett et al., 2004; Smallhorn et al., 2004), a Wnt–Paraxis–Rac1 axis may coordinate epithelialization during somitogenesis. Rac1/Cdc42 inhibition also reduced the Wnt target *Msgn1*, consistent with Rac1 influencing Wnt signaling, potentially in a reciprocal manner. Despite these perturbations, anterior–posterior polarity was largely preserved, as *Uncx4.1* expression remained confined to the posterior half of somites; qPCR reductions in *Uncx4.1* and *Meox1* likely reflect fewer, larger somites rather than disrupted patterning. Together, these findings suggest that Rac1 modulates somite morphology and epithelialization via interactions with Paraxis and Wnt signaling, with compensatory Paraxis upregulation helping maintain somite polarity despite Rac1 inhibition.

Naganathan et al. (2022) suggested that somite symmetry is established through surface tension independently of the segmentation clock, aligning with the key role of Rac1/Cdc42 in regulating cell adhesion and, consequently, somite mechanical properties. This work (Naganathan et al., 2022) indicates that Rac1 may also influence the establishment of left-right symmetry, presenting an interesting avenue for future investigation. An earlier study (Dias et al., 2014) proposed that somitogenesis can occur without the segmentation clock; however, our data show that somite formation in Rac1/Cdc42-inhibited explants remains coupled to clock periodicity, as increased oscillation period is followed by fewer somites, challenging this idea. Moreover, somites maintained *Uncx4.1* expression under perturbation, supporting a direct link between segmentation clock timing and somite formation.

In this study, we identify Rac1 as a critical regulator of oscillation dynamics in the mouse PSM, likely acting through its downstream effector PAK1. Our findings reveal general principles of cell–cell communication and pathway interactions that may extend to other tissues and morphogenetic events during embryogenesis and adult homeostasis. The Rac1–PAK–Wnt signaling axis proposed here as a key modulator of PSM oscillation dynamics warrants further investigation. Importantly, our discovery has potential clinical relevance: congenital disorders of somitogenesis, including VACTERL association and Spondylocostal Dysostosis, have long been linked to segmentation defects (Chen et al., 2016; Nóbrega et al., 2021; Sparrow et al., 2006; Turnpenny et al., 2007), and recent evidence implicates mutations in the Rac1–PAK1 pathway in VACTERL patients (Seyama et al., 2023; Solomon, 2011). By incorporating Rac1 into the molecular framework of the segmentation clock, this work opens new avenues for investigating how Rho GTPases integrate with signaling cascades and gene regulatory networks to control oscillatory periodicity—an aspect of clock regulation not previously considered. Placing Rac1 within the core framework of the segmentation clock expands the conceptual landscape of oscillatory regulation and provides a foundation for systematic exploration of Rho GTPases in developmental timing.

## Materials and Methods

### Experimental mouse models

All experiments quantifying oscillations used the LuVeLu transgenic reporter line as previously described (Aulehla et al., 2008). Male transgenic mice were time-mated with wild-type CD1 females, and embryos were collected at E9.5 for mPSMs and at E10.5 for ePSMs and tail explants. Only transgenic males were used to maintain the colony. Animals were housed at the FMI mouse facility under the license of the Basel City Veterinary Cantonal Authorities, and all experiments were conducted according to permit #2911.

### PSM reaggregation assay (ePSM)

E10.5 LuVeLu-positive embryos were screened for reporter expression and dissected to isolate PSM tissue. Four to six PSMs were pooled per experiment and mechanically dissociated by pipetting to generate single-cell suspensions (Tsiairis and Aulehla, 2016). Suspensions were filtered through 30 μm CellTrics filters, centrifuged at 400 rcf for 4 min to pellet the cells, and pellets were carefully detached using a tungsten wire. Pellets were cut into four to six pieces, and one to two pieces were transferred into each well of a fibronectin-coated 4-well chamber slide (Lab-Tek). Fibronectin coating was prepared by adding 80 μL fibronectin to 1520 μL PBS, with 360 μL applied per well for >1 h. Wells were washed and equilibrated with 320 μL DMEM-F12 (0.5 mM glucose, 2 mM glutamine, 1% BSA, 0.025/0.04 mg/mL penicillin/streptavidin) at 37°C and 5% CO₂ for ≥1 h. ePSM pieces were then transferred into the wells and imaged for ≥15 h. All steps prior to transfer were performed in medium supplemented with 10 mM HEPES.

### Chemical compound screen on the mPSMs

E9.5 LuVeLu-positive embryos were dissected to isolate posterior PSM tissue (tail bud). One to six mPSMs were plated per fibronectin-coated well of a 4-well chamber slide and equilibrated as above. Each well received a single chemical compound or a negative control (DMSO, H₂O, or methanol), matching solvent type and maximum concentration used. Compound concentrations were determined from literature; some compounds were tested at multiple concentrations. One to three compounds plus control were tested per experiment depending on sample availability. Each compound was applied in at least two independent experiments with a minimum of three replicates. mPSMs were imaged for ≥15 h at 37°C and 5% CO₂. Compound details are in Supplementary Table 1.

### Delayed and reversible chemical treatments

For delayed treatment experiments, mPSMs were treated with DMSO (0.05%) or AZA1 (2.5 μM or 5 μM) ∼6 h after imaging onset and maintained for the remainder of imaging (≥15 h total). To test reversibility, compounds were removed after ∼6 h by medium exchange with compound-free DMEM-F12, and imaging continued for ≥15 h.

### Live imaging

Confocal time-lapse imaging was performed on a ZEISS LSM 710 microscope using 514 nm excitation for LuVeLu-YFP. A 20× Plan-Apochromat objective (NA 0.8) captured three Z-planes per sample (5 μm spacing) every 10 min.

### Image processing

All images (512×512 pixels, 1.38 μm/pixel, 12-bit) were analyzed in Fiji/ImageJ (Schindelin et al., 2012). Maximum intensity projections of the Z-stacks were Gaussian-blurred (σ=10.0) and segmented using the Huang threshold. Mean fluorescence intensity and area of the segmented mPSMs were extracted over time. ePSM synchrony was assessed by drawing three non-overlapping circular regions per sample; mean fluorescence intensity per region was plotted over time.

### Time series analysis

Data were imported into R (v4.3.1) for analysis, with FDR correction applied for multiple comparisons. Period extraction used either Singular Spectrum Analysis (SSA, RSSA package) or simple cosine fitting, consistently applied within each experimental setup.

### mPSM oscillation quantification

SSA was applied in two steps: detrending (window length = 12 points) and dominant periodicity detection (window length = half the series length). Elementary time series were grouped using decomposition plots and w-correlation matrices. The first group usually represented oscillatory signal; noisy signals used the second group. Most time series were analyzed within 100–1000 min to reduce noise.

### ePSM oscillation quantification

Amplitude and period were obtained using simple cosine fits after detrending with a loess filter (span = 0.5). Amplitude was normalized to the median trend value to account for variability in fluorescence intensity.

### Area increase quantification

Sample spreading was quantified by deriving the first derivative of the area versus time plot (100–500 min) and smoothing via loess filter. The mean smoothed value represented the spreading rate.

### Analysis of delayed chemical treatments

Signals were segmented into pre-(300–30 min before treatment) and post-treatment windows (150–600 min after treatment) for period analysis via cosine fitting. Phase analysis used slightly shorter post-treatment windows (30–250 min) to account for transient peaks.

### Analysis of compound removal

Signals were segmented into pre-(290–30 min) and post-removal (30–450 min) windows, with period determined via cosine fitting.

### Measurement of somite numbers in PSM explant-cultures

E10.5 tail explants containing the last two somites were cultured in 4-well chamber slides with 320 μL medium and chemical treatment for 21.5 h (37°C, 5% CO₂, 60% O₂). Somite counts were corrected for the two pre-existing somites and normalized to culture duration.

### Quantitative PCR (qPCR)

After 22 h culture, tail explants were dissected into PSM and somite portions. RNA was extracted, quantified, and converted to cDNA (2 ng/μL). qPCR was performed with the StepOnePlus Real-Time PCR system using Platinum SYBR Green SuperMix-UDG w/ROX. Primer sequences:

- Hes5: Fw GAAGGCCGACATCCTGGAGA, Rv ACCAGGAGTAGCCCTCGCTGTA
- Dusp4: Fw AGCATGTGTGTGCAGGAGTC, Rv ACAGACCGCTGGAGAGAAAA
- Msgn1: Fw CTTCTGACACCGCTGGTCTG, Rv GTGACTGCCGTAGCCATCG
- Actin: Fw GGCTGTATTCCCCTCCATCG, Rv CCAGTTGGTAACAATGCCATGT
- Uncx4.1: Fw ACCCGCACCAACTTTACCG, Rv TGAACTCGGGACTCGACCA
- Paraxis: Fw AAGGTGCCCAGGAAGACGGGG, Rv TCATCTCCGTGCCACTCGCAG
- Meox1: Fw TGAGACGGAGAAGAAATCATCCA, Rv CGTAGCTGCTCCTTGGTGAAG

Expression levels were normalized to Actin.

### In situ hybridisation

Probes and whole-mount in situ hybridization were performed as previously described (Aulehla et al., 2008; Lauschke et al., 2013), using an *Uncx4.1* probe (Mansouri et al., 1997). Tail explants were transferred to Eppendorf tubes for processing, and littermates were processed in parallel. Images were acquired with a Leica MZ16F stereomicroscope and DFC420C camera.

### Tail and somite length measurements

Tail lengths were measured in Fiji using a 1-pixel-width segmented line along the midline in *Uncx4.1* images. Somite lengths were measured by applying a Gaussian blur and drawing a 100-pixel-width segmented line along the lateral embryo side. SSA was used to quantify intensity periodicity, corresponding to average somite length.

## Acknowledgements

We thank our lab members and the FMI community for their support and feedback. We are grateful to Maria Pappa’s PhD committee—Prof. Antoine Peters and Dr. Pierre-François Lenne—for their guidance. We also thank Zoë Gruenig, Jennifer Groeli, and the FMI animal facility staff for assistance with mouse work, as well as the FMI Imaging and Computational Facilities, particularly Drs. Jan Eglinger, Laurent Gelman, Panagiotis Papasaikas, and Dimosthenis Gaidatzis. Finally, we acknowledge the Liberali, Turco, and Grosshans labs for constructive discussions.

## Data Availability

All relevant data are included in the manuscript and are available from the corresponding author upon request.

